# Feeding Behavior Modifies the Circadian Variation in RR and QT intervals by Distinct Mechanisms in Mice

**DOI:** 10.1101/2023.11.02.565372

**Authors:** Makoto Ono, Don E. Burgess, Sidney R. Johnson, Claude S. Elayi, Karyn A. Esser, Tanya S. Seward, Carie R. Boychuk, Andrés P. Carreño, Rebecca A. Stalcup, Abhilash Prabhat, Elizabeth A. Schroder, Brian P. Delisle

## Abstract

Rhythmic feeding behavior is critical for regulating the phase and amplitude in the ≍24-hour variation of the heart rate (RR intervals), ventricular repolarization (QT intervals), and core body temperature in mice. We hypothesized the changes in cardiac electrophysiology associated with feeding behavior were secondary to changes in core body temperature. Telemetry was used to record electrocardiograms and core body temperature in mice during ad libitum-fed conditions and after inverting normal feeding behavior by restricting food access to the light cycle. Light cycle-restricted feeding quickly modified the phase and amplitude of the 24-hour rhythms in RR intervals, QT intervals, and core body temperature to realign with the new feeding time. Heart rate variability analysis and inhibiting β-adrenergic and muscarinic receptors suggested that the changes in the phase and amplitude of the 24-hour rhythms in RR intervals were secondary to changes in autonomic signaling. In contrast, the changes in the QT intervals closely mirrored changes in core body temperature. Studies at thermoneutrality confirmed the daily variation in the QT interval, but not the RR interval, and reflected daily changes in core body temperature (even in ad libitum-fed conditions). Correcting the QT interval for differences in core body temperature helped to unmask QT interval prolongation after starting light cycle-restricted feeding and in a mouse model of long QT syndrome. We conclude feeding behavior alters autonomic signaling and core body temperature to regulate the phase and amplitude in RR and QT intervals, respectively.

## Introduction

Organisms have evolved circadian rhythms in physiology to anticipate changes during the 24 hours in the external environment.(1) In mammals, circadian rhythms are generated at the cellular level by the circadian clock mechanism, a transcription-translation feedback loop that cycles with a periodicity of ≈24 hours.(2, 3) The circadian clock mechanism in the suprachiasmatic nucleus (SCN) of the hypothalamus entrains to the light cycle, and light-entrained SCN neurons signal to neighboring SCN neurons and other brain nuclei to align neurohumoral signaling, behavioral rhythms, and peripheral circadian clocks throughout the body.(4, 5) Several studies suggest SCN signaling modifies autonomic input to the heart to generate circadian variations in cardiac electrophysiology (i.e., RR and QT intervals).(6, 7) Studies also show that the circadian clock mechanism in cardiomyocytes contributes to 24-hour rhythms in the RR and QT intervals by altering the expression of genes, proteins, and different cardiac ionic currents.(8-14)

Feeding behavior is an important timekeeping cue for aligning circadian clocks in peripheral tissues, including the heart. Mouse studies reveal that restricting feeding to the inverse time of normal feeding behavior (i.e., light cycle-restricted feeding) realigns the phase of circadian mRNA expression in the heart and 24-hour rhythms in cardiac electrophysiology to the new feeding behavior.(15, 16) The effect of light cycle-restricted feeding on the realignment of 24-hour rhythms in cardiac electrophysiology is not secondary to changes in the circadian clocks in the SCN or cardiomyocytes. The reason is the circadian clock in the SCN remains entrained to the light cycle during light cycle-restricted feeding(17), and genetic disruption of the cardiomyocyte circadian clock by inducing deletion of BMAL1 does not prevent realignment of 24-hour rhythms in cardiac electrophysiology with feeding.(18) This study determined how feeding behavior impacts 24-hour rhythms in the phase and amplitude of the RR and QT intervals in mice.

Several studies have demonstrated a strong temperature dependence in the RR and QT interval.(19-29) In addition, previous studies have shown that light cycle-restricted feeding in mice alters the phase and amplitude of the 24-hour rhythm in core body temperature.(17, 30) This raises the intriguing possibility that feeding modifies 24-hour cardiac electrophysiology rhythms by altering core body temperature rhythms.

In this study, we tested whether light-cycle restricted feeding similarly altered the phase and amplitudes of the 24-hour rhythms of the RR intervals, QT intervals, and core body temperature. We found that the changes in the 24-hour rhythm of the RR interval during light cycle-restricted feeding were only partially explained by changes in the core body temperature, and pharmacological studies showed the 24-hour rhythm in the RR interval was strongly influenced by autonomic signaling. In contrast, the changes in the 24-hour rhythm in the QT interval during light cycle-restricted feeding closely mirrored changes in the core body temperature. Correcting the QT interval for core body temperature was important in identifying abnormal changes in the QT interval in transgenic mouse models of long QT syndrome. We conclude that feeding behavior aligns the 24-hour rhythms in the RR and QT intervals by distinct mechanisms in mice.

## Methods

### Animals

The mice used in this study were male on the 129S6 background strain. The 129S6 mouse strain has been previously characterized in the ΔKPQ mouse model (*Scn5a^+/^*^Δ*KPQ*^ mice).(31, 32) For studies comparing the genotype-specific differences, age-matched (wild type, WT) and *Scn5a^+/^*^Δ*KPQ*^ littermates were used. Mice were fed Teklad Global 18% protein rodent diet (2018). All procedures complied with the Association for Assessment and Accreditation of Laboratory Animal Care guidelines and were approved by the Institutional Animal Care and Use Committee at the University of Kentucky. Dr. Peter Carmeliet kindly provided the *Scn5a*^+/ΔKPQ^ mouse strain.(33) This study represents new and re-analysis of data collected previously.(15) All studies were performed at room temperature (23 ± 2^°^C) except for data presented in **Figure 7**. These studies were done in mice housed in thermoneutral temperatures (30 ± 2^°^C).(34) After the study, mice were euthanized by anesthetic overdose followed by cervical dislocation.

### Telemetry

We used telemetry to measure electrocardiogram (ECG), core body temperature, and locomotor activity. Mice were anesthetized with isoflurane (2-4%) using a nose cone attached to an anesthetic gas vaporizer and implanted with PhysioTel ETA-F10 (Data Sciences International) telemetry transmitter units. ECG telemetry data were recorded at 1000 Hz, and RR, QT, and PR interval analyses were done using MATLAB (MathWorks), similar to those previously described.(8, 15, 18) ECG traces recorded for each hour were aligned to the peak of the R wave to generate an ensemble ECG trace. The beginning of the P wave was defined as the point where the slope of the P wave profile changed by more than 0.001 mV/ms; the Q wave was defined as the point where the slope of the profile changed from negative to positive at the base of the QRS complex; the end of the T wave was defined as the point of intersection of T wave maximum slope with baseline.(35)

### Statistical analysis

For some graphs, data are shown as the mean and standard deviation of the mean (SD). For other graphs, a data point for each animal is shown with a horizontal bar at the mean. The statistical JTK_CYCLE package confirmed significant cyclical rhythms at p<0.05 before analyzing the data using cosine waveform curve fitting.(36). Data were analyzed in PRISM (GraphPad). Details of statistical analyses for each experiment are reported in the figure legends.

## Results

### Experiment 1: Determine how long it takes for light cycle-restricted feeding to alter the 24-hour rhythms in cardiac electrophysiology and core body temperature

Three to four-month-old WT mice housed in 12-hour light/dark cycles at room temperature with ad libitum access to water were used in this study. To initiate light cycle-restricted feeding, food was removed at 9 h after the beginning of the light cycle (Zeitgeber time or ZT = 9). The next day, food presentation was restricted to 7 h during the light cycle, between ZT 2–9 for two weeks. Light cycle-restricted feeding caused mice to lose body mass during the first week (**Figure S1**). However, caloric intake increased after that, and similar to previous reports, light cycle-restricted-fed animals re-gained body mass after two weeks.(37)

We used *in vivo* telemetry to longitudinally record electrocardiogram (ECG) data from mice before and after switching to light cycle-restricted feeding (**Figure 1A**). Data recorded shortly after starting light cycle-restricted feeding revealed immediate impacts on cardiac electrophysiology. Data recorded two weeks after beginning light cycle-restricted feeding indicated the steady state changes.(17) The top, left panel in **Figure 1B** shows a three-day time series for changes in the hourly mean heart rate (RR interval), the duration of ventricular repolarization (QT interval), and the duration of atrial to ventricular conduction (PR interval) in ad libitum-fed conditions. The time series showed that the amplitude in the daily rhythms in RR, QT, and PR intervals peaked at a similar time (i.e., had similar acrophases) during the light cycle. However, a new 24-hour rhythm in the ECG data emerged one day after starting light cycle-restricted feeding. The new 24-hour rhythms had larger amplitudes and peaked in the dark cycle. The changed 24-hour rhythms in the ECG data after starting light cycle-restricted feeding were not transitory and persisted for two weeks.

**Figure 1.**
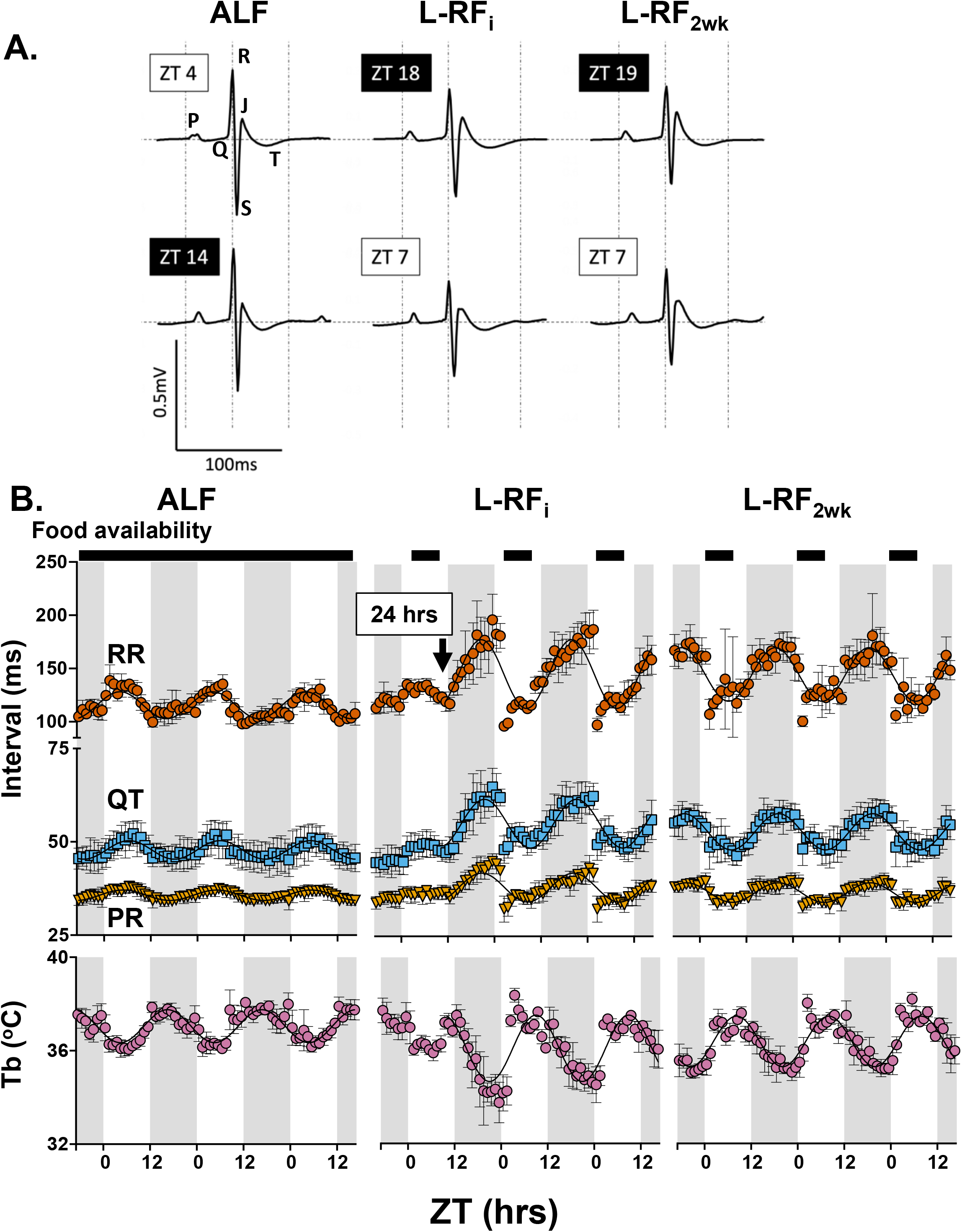
Light cycle-restricted feeding caused large changes in hourly RR intervals, QT intervals, PR intervals, and core body temperature. A. Representative ensemble ECG traces recorded from a mouse in ad libitum-fed conditions (ALF, left column), after initially starting light cycle-restricted feeding (L-RF_i_, middle column), and two weeks after starting light cycle-restricted feeding (L-RF_2wk_, right column) at different Zeitgeber times (ZT). B. Averaged time series data showing three consecutive days of the hourly mean for the RR interval (vermillion), QT interval (sky blue), PR interval (orange), and core body temperature (Tb, reddish purple). The data were plotted as a function of ZT and the shaded regions correspond to the dark cycle. The left column shows the data recorded during ALF, the middle column shows the data recorded during L-RF_i_ (the boxed arrow indicates 24 hours after the start of light cycle-restricted feeding), and the right column shows the data recorded after L-RF_2wk_. The solid lines are cosine curves fitted to the data (n = 6 mice, data points are averaged data and SD).

Previous studies show that light cycle-restricted feeding causes analogous changes in daily core body temperature rhythms.(17, 38) We found that light cycle-restricted feeding changed the daily core body temperature rhythm in a way that largely mirrored the changes in the daily rhythms in RR, QT, and PR intervals (**Figure 1B**).

We quantified the changes in the 24-hour rhythms in cardiac electrophysiology and core body temperature before and after starting light cycle-restricted feeding. Almost all of the individual ECG and core body temperature time series recorded from ad libitum-fed mice, one day after starting light cycle-restricted feeding, and two weeks after starting light cycle-restricted feeding had significant cyclical rhythms with a periodicity of ≍24 hours (i.e, JTK analysis showed p<0.05) (see **Supplemental Table 1**).

The hourly mean data for RR, QT, and PR intervals and core body temperature for each mouse were fit with the cosine curve function:

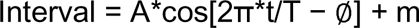

to calculate the period (T), the time between the peak amplitudes; acrophase (∅), the phase of the peak amplitude in relation to the start of the light cycle (ZT = 0); the amplitude (A), one-half of the peak to trough; and the mesor (m), the rhythm adjusted mean (calculated data are presented in **Supplement Table 1**).(8, 15, 18)

Data showed that light cycle-restricted feeding caused large changes in the acrophase, amplitude, and mesor of daily rhythms in RR intervals, QT intervals, PR intervals, and core body temperature (**Figures 2A, 2B**). No or very small additional changes were seen in the cyclical ECG and core body temperature rhythms two weeks after starting light cycle-restricted feeding.

**Figure 2.**
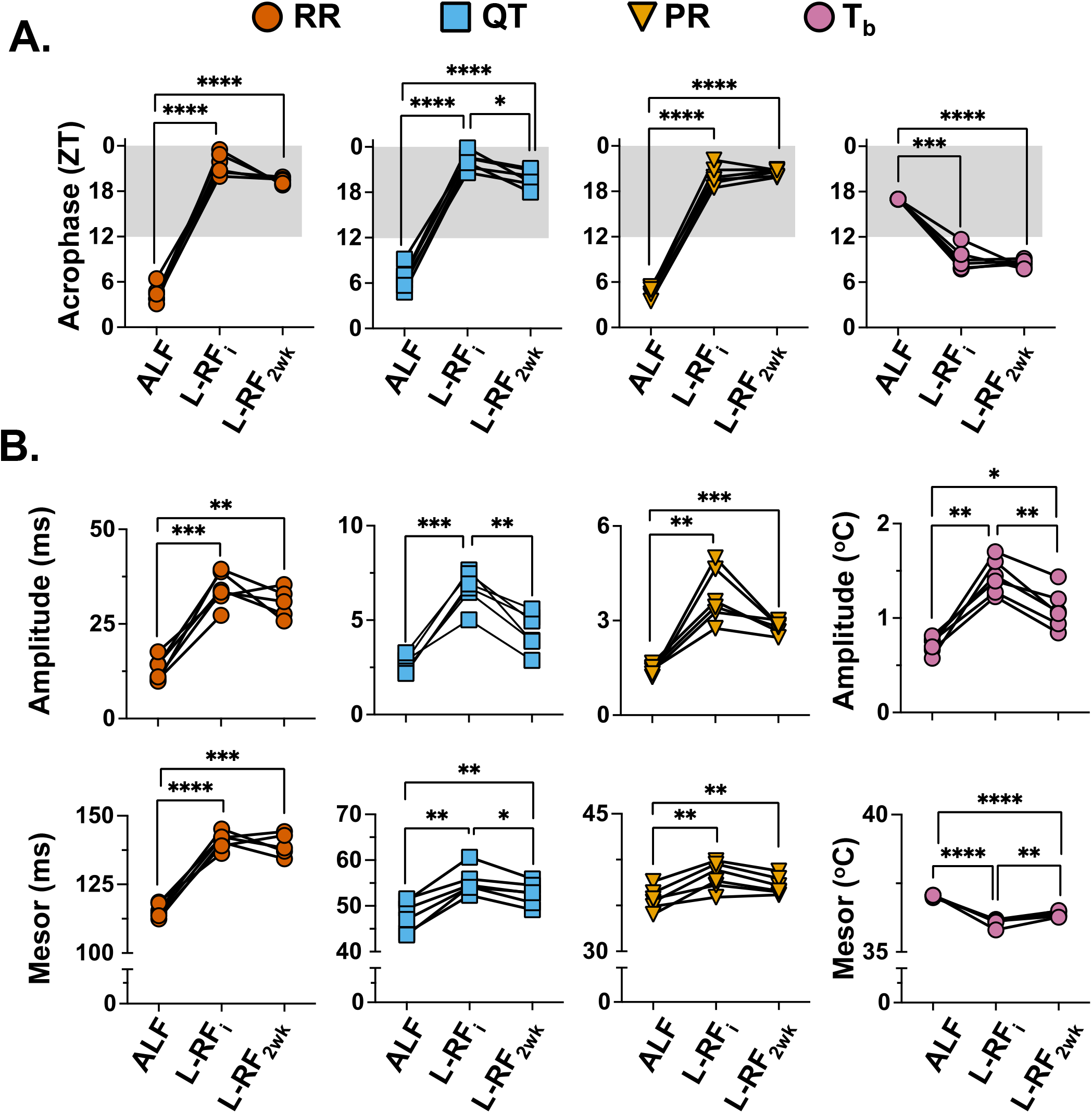
Light cycle-restricted feeding modified the properties of the 24-hour rhythms for RR intervals, QT intervals, PR intervals, and core body temperature. The individual mouse time series data for the RR intervals, QT intervals, PR intervals, and core body temperature (Tb) were fit to a cosine wave to calculate the acrophase, amplitude, and mesor for mice in ad libitum-fed conditions (ALF), 24 hours after initially starting light cycle-restricted feeding (L-RF_i_), and two weeks after starting light cycle-restricted feeding (L-RF_2wk_). **See Supplemental Table 1. A.** The top row shows the acrophase of the RR interval (vermillion), QT interval (sky blue), PR interval (orange), and Tb (reddish purple) for each condition. **B.** The top row shows the amplitude, and the bottom row shows the mesor of the RR interval, QT interval, PR interval, and Tb for each condition. Data were analyzed using a two-way mixed effects model (**see Figure 3S**) with Geisser and Greenhouse correction and Sidǎk correction, *p<0.05, **p<0.01, ***p<0.001, ****p<0.0001, n = 5-6 mice per group.

We conclude only one day of light cycle-restricted feeding causes large, simultaneous, and abrupt changes in the phase, amplitude, and mesor in the ≈24-hour rhythms of cardiac electrophysiology and core body temperature.

### Experiment 2: Light cycle-restricted feeding impacts the 24-hour rhythm in the RR interval by modifying core body temperature and autonomic input to the heart

We first estimated how much of the 24-hour rhythm in the RR interval is expected to be driven by daily change in core body temperature (RR_core body temperature_ interval). We used a Q_10_ value based on the Q_10_ value reported in isolated WT mouse hearts.(23) For the RR_core body temperature_ interval calculation, we used a reference core body temperature of 37^°^C and an RR interval of 116 ms because these were the mean values in WT mice in ad libitum-fed conditions.

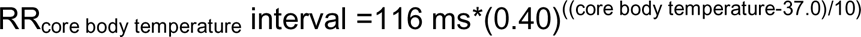

We directly compared the estimated RR_core body temperature_ interval with the RR interval measured from WT mice in ad libitum-fed conditions, immediately after starting light cycle-restricted feeding, and two weeks after starting light cycle-restricted feeding (**Figure 3**). The data suggest that the RR_core body temperature_ interval represented a significant portion of the amplitude in the RR interval during ad libitum-fed conditions. However, a large difference between the RR_core body temperature_ and RR interval emerged during the dark cycle one day after starting light cycle-restricted feeding (**Figure 3A**), suggesting that additional factors contributed to the amplitude of the 24-hour rhythm in RR intervals one day after starting light cycle-restricted feeding.

**Figure 3.**
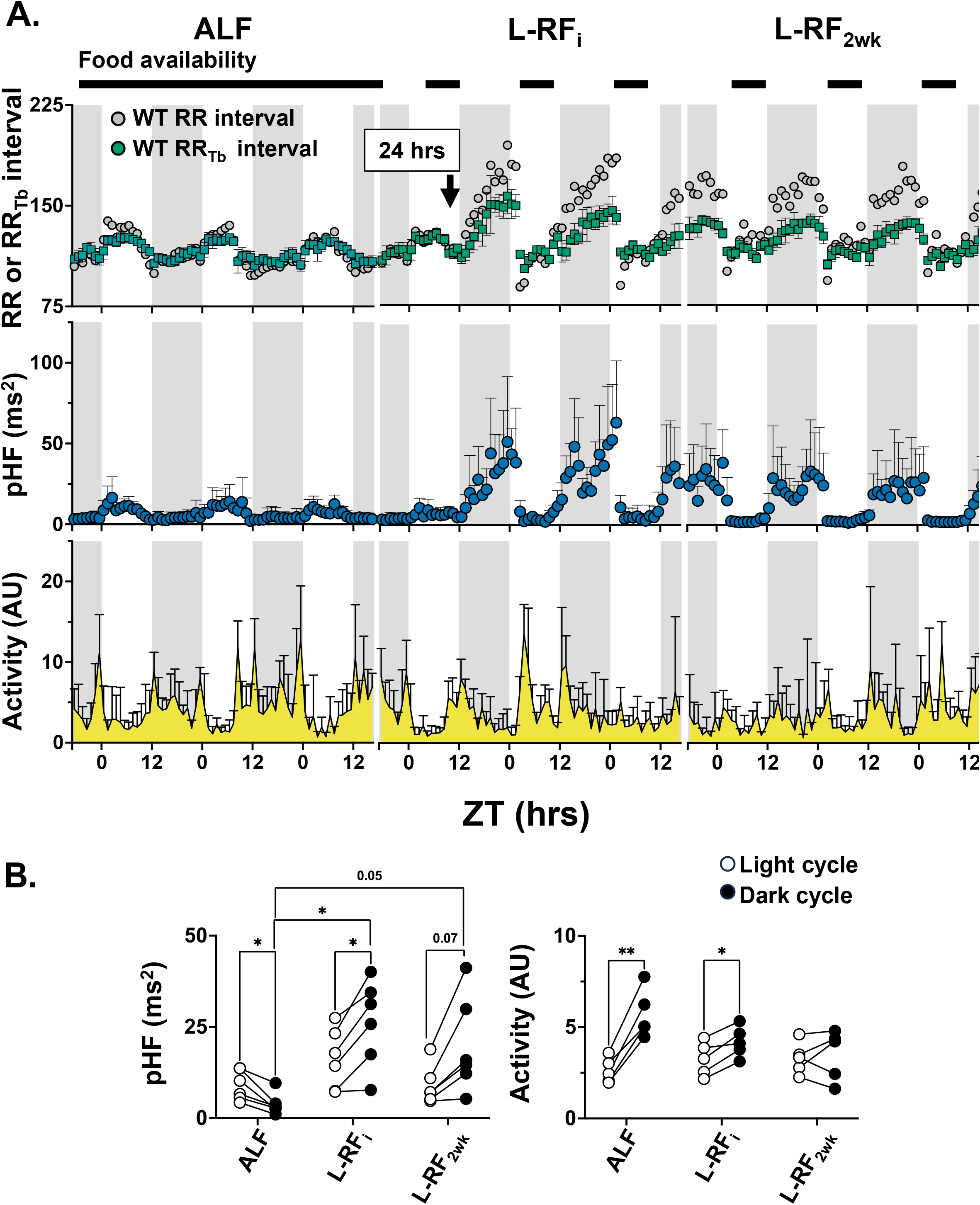
Light cycle-restricted feeding altered heart rate variability and activity. **A**. The top row of graphs shows the averaged time series data for three consecutive days of the hourly mean for the RR interval from **Figure 1** (grey) overlayed with the estimated RR_core body temperature_ interval (green). The left column shows the data from mice in ad libitum-conditions (ALF), the middle column shows the data initially starting light cycle-restricted feeding (L-RF_i_), and the third column shows the data two weeks after starting light cycle-restricted feeding (L-RF_2w__k_). The middle row of graphs shows the corresponding data for the power of the high-frequency component in heart rate variability (pHF, blue), and the bottom row of graphs shows the corresponding data for cage activity (yellow). Data were plotted as a function of ZT and the shaded regions correspond to the dark cycle (n = 6 mice, data points are averaged data and SD). **B.** The left graph shows the average pHF value for each mouse during the light cycle (open circle) and dark cycle (solid circle) measured during ALF, L-RF_i_, and L-RF_2wk_. The right graph shows the average activity value for each mouse during the light cycle (open circle) and dark cycle (solid circle) during ALF, L-RF_i_, and L-RF_2wk_. Data were analyzed using a one-way ANOVA with Geisser and Greenhouse correction and Sidǎk correction, *p<0.05 and **p<0.01, n = 6 mice per group.

The 24-hour rhythm in the RR interval is classically thought to reflect changes in autonomic input to the heart and activity.(6, 39, 40) We quantified changes in autonomic input to the heart by measuring changes in the heart rate variability from mice in ad libitum-fed conditions and after switching to light cycle-restricted feeding. We followed the approaches outlined by Fenske et al. (41). Interpolated RR interval time series were generated for spectral analysis and determination of frequency domain components for heart rate variability analyses. In mice, the frequency range to classify the high-frequency (HF) component is 1.5-4.0 Hz, and the frequency range used to classify the low-frequency (LF) component is 0.4-1.5 Hz (41) The HF component is generally accepted to correspond to the parasympathetic outflow related to respiratory sinus arrhythmia, and the LF component is thought to primarily reflect autonomic outflow associated with the baroreceptor reflex.(42)

Power Spectral Density *𝒫*(*f*) has units of power per frequency (ms^2^/Hz). The HF and LF components represent the power within a frequency band and are related to the integration:

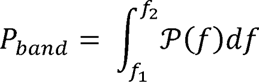

The units of the differential cancel the frequency units, so the units for the HF and LF components are in ms^2^. Normalization is based on the discrete version of Parseval’s theorem. It allows us to consistently approximate the integration for LF and HF components of the variance for the underlying RR time series.(43)

In ad libitum-fed conditions, the mean HF component was higher during the light cycle than the dark cycle. One day after starting light cycle-restricted feeding, the mean HF component increased during the dark cycle, and this persisted two weeks after starting light cycle-restricted feeding (**Figure 3A, 3B**). Similar trends were seen for the LF component but did not reach statistical significance (**Figure 3S**).

In ad libitum-fed conditions, the mean activity was lower during the light cycle than the dark cycle. This trend persisted initially after starting light cycle-restricted feeding, but the difference between the light and dark cycle activity levels disappeared after two weeks.

The individual mouse three-day time series data for the HF and activity components did not show significant cyclical rhythms for most mice (JTK analysis P>0.05), and the data were not well described using cosine waveform curve fitting. Therefore, we analyzed the relative rhythmicity between the RR interval and the HF component or activity using cross-correlation analyses.(44) As part of the analysis, we included the cross-correlation between RR interval and core body temperature. We used the R function “ccf” to calculate the cross-correlation.(45) The formula used in “ccf” to calculate the cross-correlation between two time series *X(t)*, *Y(t)* has the form

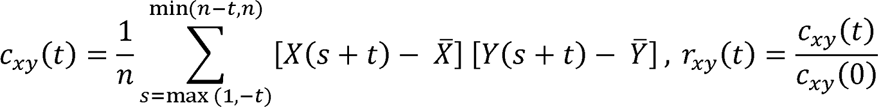

Figure 4 shows the cross-correlation for RR-core body temperature, RR-HF, and RR-activity from mice in ad libitum-fed conditions and after starting light cycle-restricted feeding. The cross-correlation between the RR-core body temperature and RR-HF showed strongly correlated rhythmicity that peaked at a lag of 0 in all three conditions, but the RR-activity cross-correlation only showed correlated rhythmicity in ad libitum-fed conditions. The data suggested that altered autonomic signaling and not activity may be important for modifying the daily rhythm in the RR interval during light cycle-restricted feeding.

**Figure 4.**
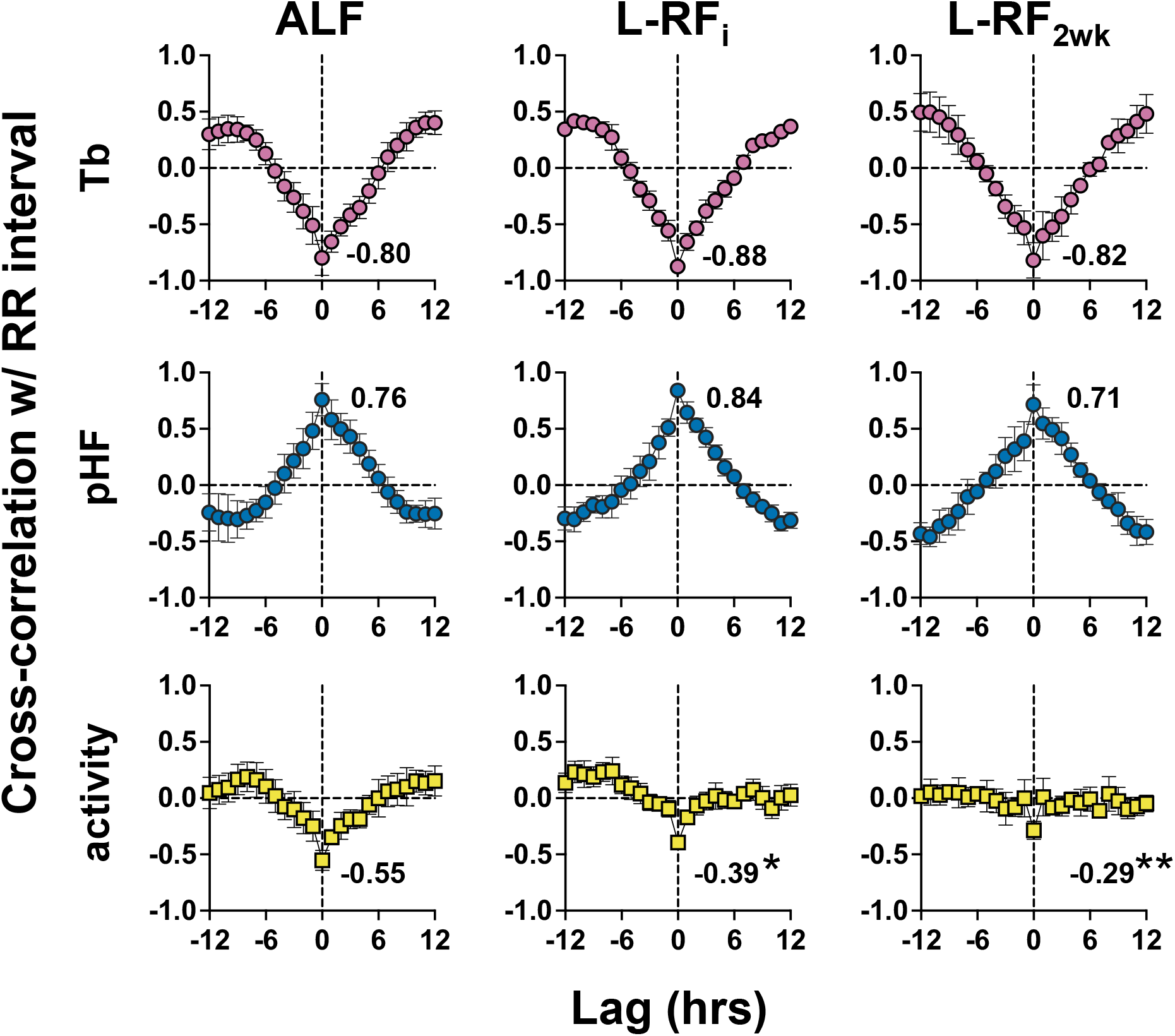
After starting light cycle-restricted feeding, the RR interval continued to show correlated rhythmicity with core body temperature and pHF but not activity. The mean data and SD for the cross-correlation between the time series data for the RR interval and core body temperature (Tb, top row, reddish purple), power of the high-frequency component in heart rate variability (pHF, middle row, blue), and activity (bottom row, yellow) in ad libitum-fed conditions (ALF, left column), after initially starting light cycle-restricted feeding (L-RF_i_, middle column), and two weeks after starting light cycle-restricted feeding (L-RF_2wk_, right column). The peak correlation coefficient at lag 0 is shown (n = 6 mice per group). Differences in the peak correlation coefficient data at lag 0 were analyzed using a one-way ANOVA with Geisser and Greenhouse correction and Sidǎk correction *p<0.05 and **p<0.01.

We directly tested the involvement of autonomic signaling by doing intraperitoneal injections of propranolol hydrochloride and atropine sulfate (10 mg/kg each) in six to eight-month-old telemetered WT mice undergoing light cycle-restricted feeding.(46-48) Propranolol and atropine inhibit β-adrenergic and muscarinic receptor activation, respectively. We injected the mice at ZT 23.5 (when the daily rhythm in the RR interval peaked) and 9.5 (when the RR interval was at its nadir) for two consecutive days. The feeding onset was delayed one hour to ZT 3 to separate the food anticipatory decrease in RR interval from the start of the light cycle. Propranolol and atropine injections caused a loss in the 24-hour rhythms of the RR interval, core body temperature, and the difference between the RR_core body temperature_ interval and RR interval (Figure 5A**, 5B, and 5C**).

**Figure 5.**
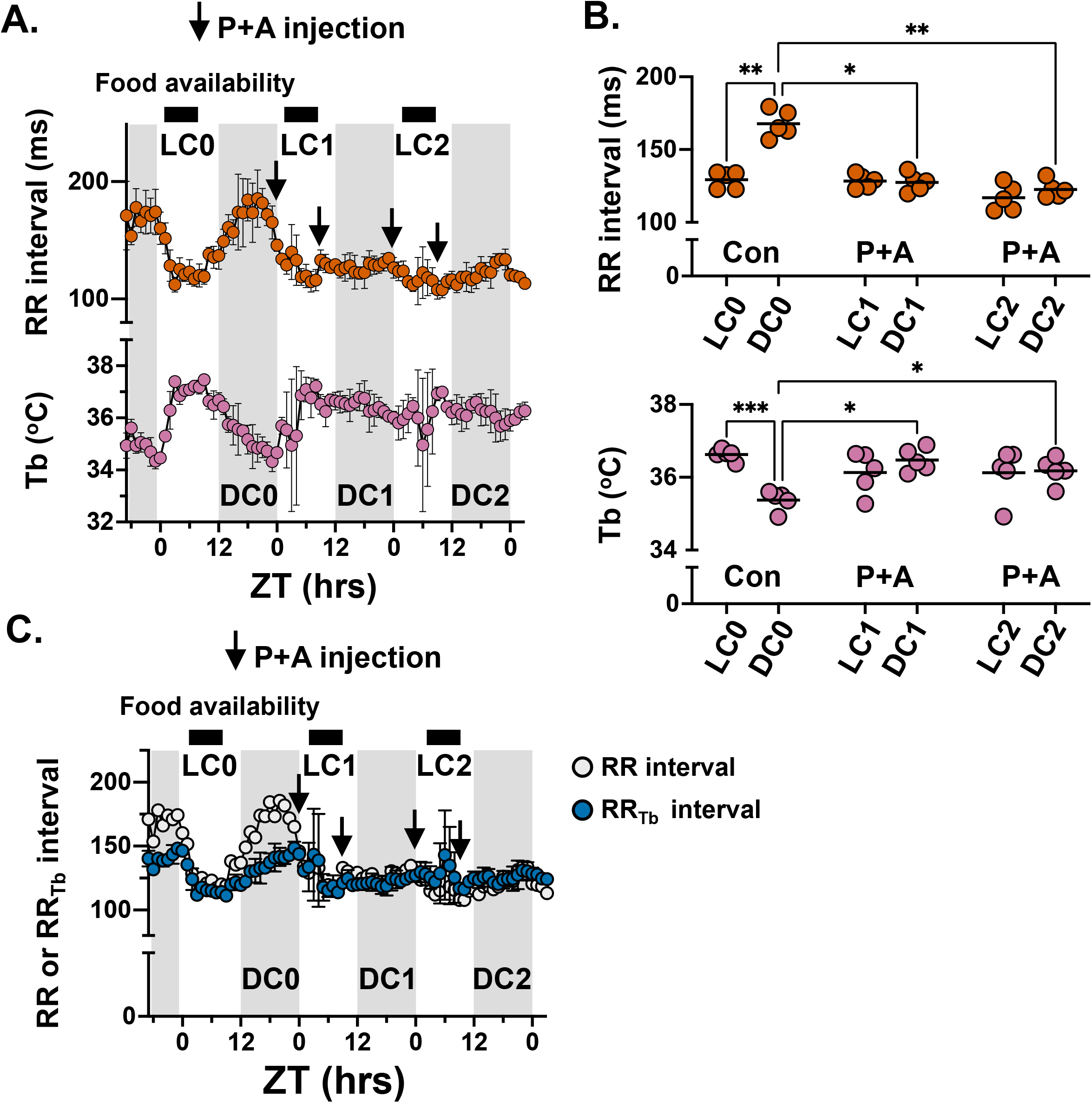
Inhibition of autonomic receptor activation caused a loss in the daily rhythm of the RR interval during light cycle-restricted feeding. A. Time series data showing the hourly mean in the RR interval (vermillion) and core body temperature (Tb, reddish purple) recorded 1 week after the start of light cycle-restricted feeding (ZT 3 to ZT 9). The data were plotted as a function of ZT and the shaded regions represent the dark cycle. The time series shows the hourly mean data and SD measured during the light and dark cycle the day before starting the propranolol and atropine injections (LC0, DC0), and the first and second light and dark cycles after starting the propranolol and atropine injections at ZT 23.5 and ZT 9.5 (LC1, DC1, LC2, and DC2, respectively, injection times are denoted by arrows; n = 5 mice). **B.** Shown are the RR interval (vermillion) and Tb (reddish purple) data recorded during LC0 and DC0, LC1 and DC1, and LC2 and DC2; the horizontal bar in each data set is the mean data. The data were analyzed using a two-way ANOVA with Geisser and Greenhouse correction and Tukey correction, *p<0.05, **p<0.01, ***p<0.001. **C.** Time series of RR interval data from **A.** overlayed with the averaged time series for the corresponding RR_core body temperature_ interval (blue) before and after injection with propranolol and atropine.

These data suggest light cycle-restricted feeding modifies the 24-hour rhythm in the RR interval by altering core body temperature and autonomic input to the heart.

### Experiment 3: The 24-hour rhythm in the QT interval during light cycle-restricted feeding primarily reflects changes in the core body temperature

The time series data shows that longitudinal changes in the hourly mean QT interval occur in parallel to the changes in the RR interval and mirror changes in core body temperature (Figure 1). The duration of the QT interval in mice is thought to be independent of the RR interval.(9, 49-51) Therefore, we quantified the relation between the QT interval and core body temperature. We used an approach similar to that described previously to correct the heart rate corrected QT interval to core body temperature in beagles.(29) The hourly mean QT intervals were plotted against the corresponding core body temperature for all data collected in ad libitum- and light cycle-restricted fed mice, and we identified a strong negative linear relationship between the mean QT interval and core body temperature (Figure 6A).(29) The average slope of the linear QT-core body temperature relation was used to derive a QT-core body temperature correction formula. We corrected the QT intervals to 37^°^C (QT_37°C_ interval) because 37^°^C was the daily mean in the core body temperature for ad libitum-fed mice.

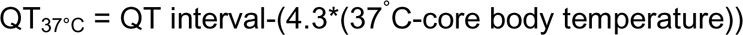

**Figure 6.**
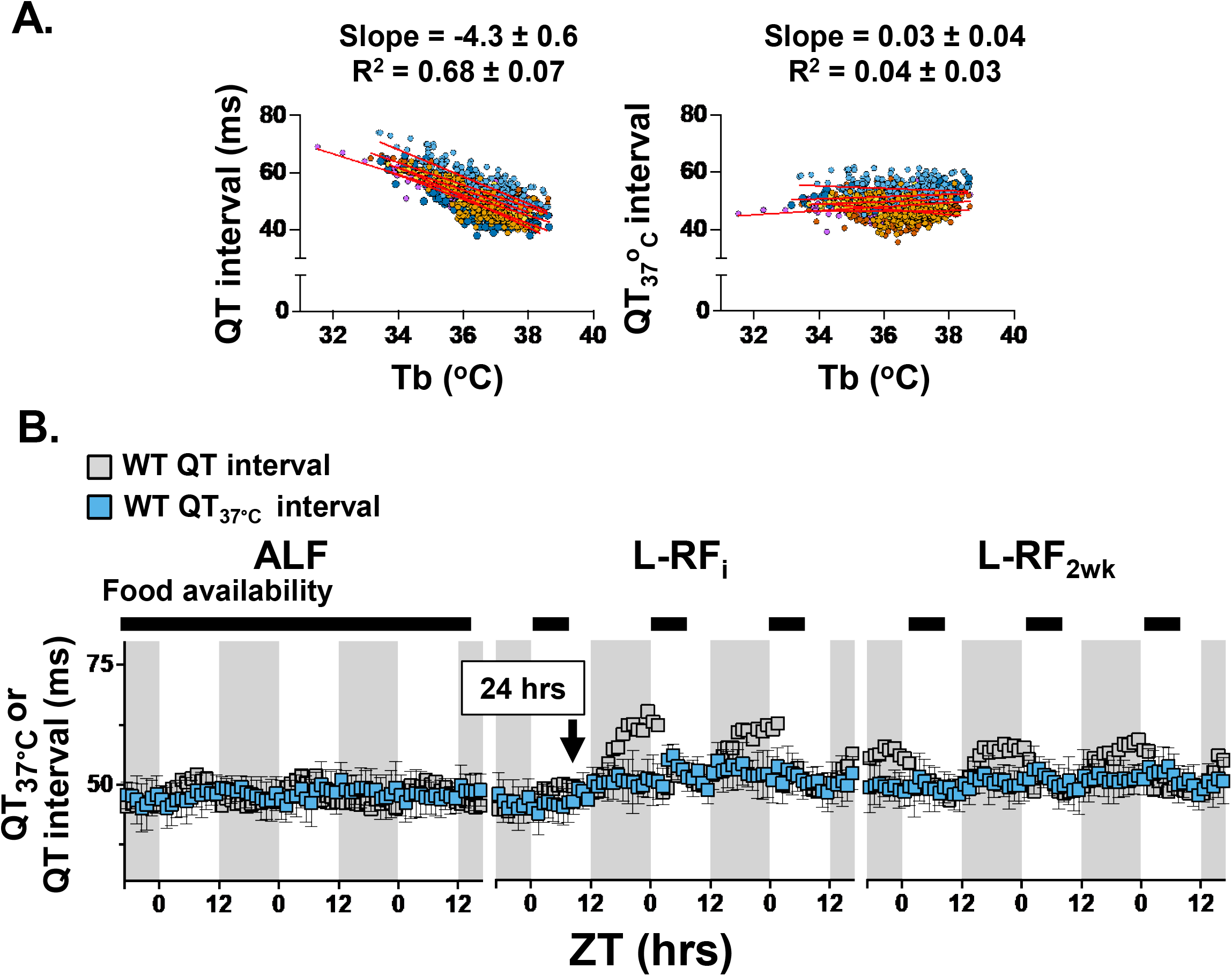
Correcting the QT interval for core body temperature caused a loss in the daily rhythm in the QT interval. A. The left graph shows the hourly mean in the QT interval plotted as a function of the corresponding core body temperature (Tb) for each mouse in ad libitum-fed conditions, initially after starting light cycle-restricted feeding, and two weeks after starting light cycle-restricted feeding (each mouse is represented by a different color, n = 6). The average slope was used to correct the QT interval to 37°C (QT_37°C_ interval). The right graph shows the QT_37°C_ interval is plotted as a function of the corresponding Tb for the same set of mice. Linear regression of the data for each mouse (red lines) was used to calculate the average slope and R^2^ and SD for both the left and right graphs. **B.** The left column shows the hourly QT_37°C_ interval and SD calculated during ad libitum-fed conditions (ALF), the middle column shows the QT_37°C_ interval initially starting light cycle-restricted feeding (L-RF_i_), and the third column shows the QT_37°C_ interval two weeks after starting light cycle-restricted feeding (L-RF_2wk_). Also shown is the corresponding time series data for the hourly mean in the uncorrected QT interval data from **Figure 1** for direct comparison (grey). Data were plotted as a function of ZT and the shaded regions represent the dark cycle.

Correcting the time series of the 24-hour rhythm in the QT interval for core body temperature caused a loss in the 24-hour rhythm in the QT interval for mice in ad libitum and light cycle-restricted feeding (Figure 6B).

We confirmed the 24-hour rhythm in the QT interval reflected changes in core body temperature (and not the RR interval) by studying the ECG properties of the mice housed in thermoneutrality under ad libitum-fed conditions. Studying mice in thermoneutrality alters autonomic tone to dramatically slow the basal heart rate without causing large changes in the daily rhythm of the core body temperature.(52) Housing mice at thermoneutrality slowed the RR interval and generated a large difference in the RR_core body temperature_ not seen in ad libitum-fed mice housed at room temperature (compare Figure 3**, top left panel** to Figure 7A). In contrast, both the QT and QT_37°C_ intervals measured in thermoneutrality were qualitatively similar to the QT and QT_37°C_ intervals measured at room temperature (compare Figure 6B, **left panel** to Figure 7A). The QT_37°C_ interval in mice housed at thermoneutrality also generated a horizontal line when plotted as a function of core body temperature (Figure 7B). We plotted the hourly mean QT intervals against the corresponding core body temperature. The negative linear relationship between the mean QT interval and core body temperature was similar to mice housed at room temperature (compare Figure 6A **and 7C**).(29)

**Figure 7.**
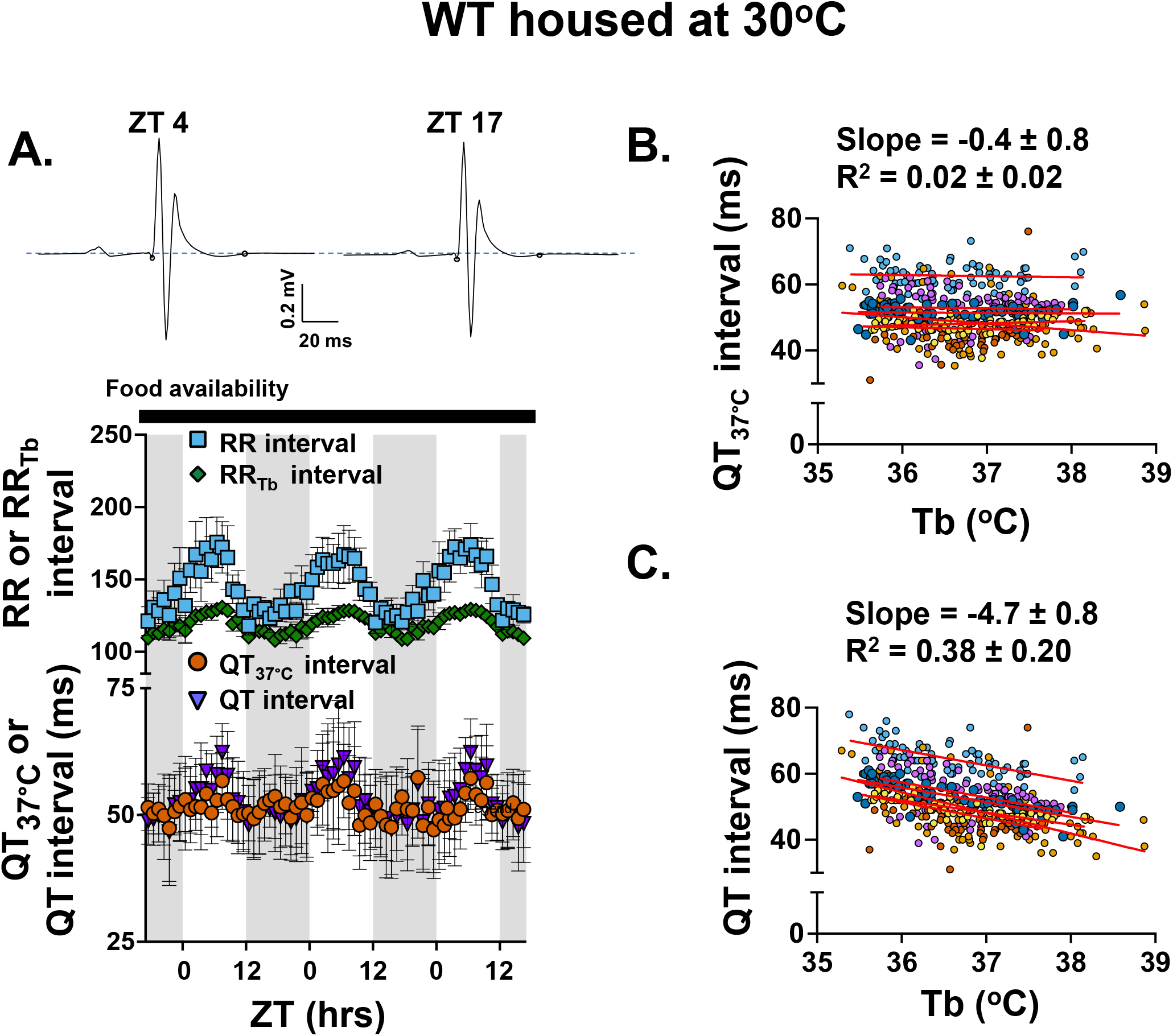
Housing mice in thermoneutrality unmasked distinct mechanisms regulating daily rhythms in RR and QT intervals. A. Representative hourly ensemble ECG traces recorded from a mouse in thermoneutrality at ZT 4 and 17. Also shown are the time series data for the hourly mean (and SD) for the RR (sky blue), RR_core body temperature_ (blue), QT (purple), and QT_37°C_ (vermillion) intervals measured from mice in ad libitum-fed conditions housed at thermoneutrality. Data were plotted as a function of ZT and the shaded regions correspond to the dark cycle (n = 6 mice). **B.** The top graph plots the QT_37°C_ interval as a function of the corresponding core body temperature (Tb) for the same set of mice in **A**. Each mouse is represented by a different color. The bottom graph shows the hourly mean in the uncorrected QT interval plotted as a function of the corresponding Tb. Linear regression (red lines) analysis for each mouse was used to calculate the mean slope and R^2^ (including SD) for both data sets.

These data support the concept that the 24-hour rhythms in the RR and QT intervals are regulated by distinct mechanisms-even in ad libitum fed mice. Specifically, the 24-hour rhythm of the RR interval is sensitive to changes in core body temperature and autonomic signaling, whereas the 24-hour rhythm of the QT interval primarily reflects changes in the core body temperature.

### Experiment 4: Determine the impact of core body temperature on the QT interval in a transgenic mouse model of long QT syndrome

We previously showed that light cycle-restricted feeding for two weeks prolonged the RR and QT intervals in *Scn5a*^+/ΔKPQ^ mice. However, the QT intervals in this study were not corrected for core body temperature.(15) We quantified the ECG properties and core body temperature measured in *Scn5a*^+/ΔKPQ^ mice during ad libitum-fed conditions and after starting light cycle-restricted feeding. The changes in the 24-hour rhythm of RR interval, QT interval, PR interval, and core body temperature were similar to those recorded in WT mice (Figures 1 **and 3S**). Cosine fitting analysis of ECG data from the *Scn5a*^+/ΔKPQ^ mice showed that, one day after starting light cycle-restricted feeding, there was an immediate change in the phase, increase in amplitude, and change in the mesor for 24-hour rhythms in the RR intervals, QT intervals, PR intervals, and core body temperature (Figure 4S). We compared the QT_37°C_ interval for WT and *Scn5a*^+/ΔKPQ^ mice in ad libitum-fed conditions and after starting light cycle-restricted feeding (Figure 8A). The QT_37°C_ interval was not different between WT and *Scn5a*^+/ΔKPQ^ mice in ad libitum-fed conditions (Figure 8B). After starting light cycle-restricted feeding, the QT_37°C_ interval became longer in WT and *Scn5a*^+/ΔKPQ^ mice. Light cycle-restricted feeding also unmasked a genotype-specific difference in QT_37°C_ intervals measured from WT and *Scn5a*^+/ΔKPQ^ mice (Figure 8B).

**Figure 8.**
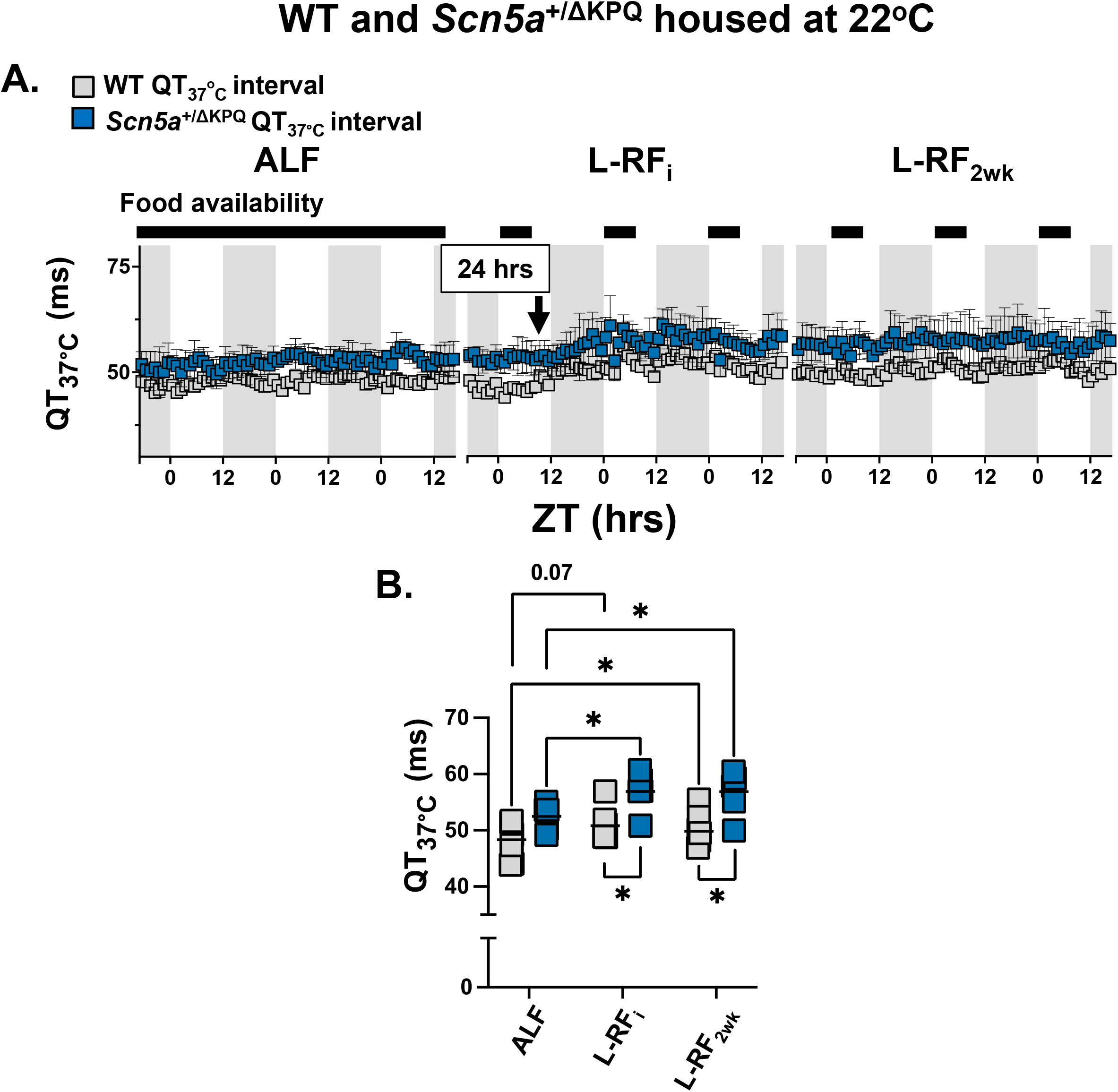
QT interval correction for core body temperature unmasked QT interval prolongation in a mouse model of long QT syndrome. **A.** The left column shows the hourly QT_37°C_ interval and SD calculated from *Scn5a*^+/ΔKPQ^ mice (blue) during ad libitum-fed conditions (ALF), the middle column shows the QT_37°C_ interval measured initially after starting light cycle-restricted feeding (L-RF_i_), and the third column shows the QT_37°C_ interval measured two weeks after starting light cycle-restricted feeding (L-RF_2wk_). The graph also shows the mean hourly QT_37°C_ interval measured from WT mice from Figure 6 for comparison (grey) C. The graph shows QT_37°C_ intervals measured from WT (grey) or *Scn5a*^+/ΔKPQ^ mice (blue) in ALF, L-RF_i_, and L-RF_2wk_. The horizontal bar in each data set is the mean (Data were analyzed using a two-way ANOVA with Geisser and Greenhouse correction and Sidǎk correction, n = 6, *p<0.05,).

## Discussion

We determined the mechanisms that underlie the changes in the 24-hour rhythms of the RR and QT intervals during light cycle-restricted feeding. We found that one day after starting light cycle-restricted feeding, the 24-hour rhythms in the RR intervals, QT intervals, PR intervals, and core body temperature changed to a new phase with a larger amplitude (Figures 1B**, 2A, 2B**). The Changes in the 24-hour rhythm of the RR interval correlated with changes in heart rate variability (Figures 3**, 4**), and inhibiting β-adrenergic and muscarinic receptor activation in light cycle restricted-fed mice caused a loss in the 24-hour rhythm of the RR interval (Figure 5). The 24-hour rhythm in the QT interval showed a strong linear relation with core body temperature (Figure 6), and studies in ad libitum-fed mice housed in thermoneutrality confirmed that the 24-hour rhythm in the QT interval primarily reflected changes in core body temperature (not the RR interval) (Figure 7). Correcting the QT interval for core body temperature allowed us to identify abnormal QT interval prolongation in a transgenic mouse model of long QT syndrome (Figure 8).

### Feeding behavior impacts the 24-hour rhythm in the RR interval (heart rate) by modifying autonomic input to the heart

Studies suggested that the circadian clock in the SCN modified autonomic signaling to generate the 24-hour rhythm in heart rate.(6, 53-56) These studies were supported by data showing that ablation of the SCN or transgenic mice lacking the core circadian clock transcription factor *Bmal1* (e.g., *Bmal1*^−/−^) did not have a 24-hour rhythm in heart rate.(57-59) Light cycle-restricted feeding is not thought to impact the phase of the circadian clock mechanism in the SCN;(17) therefore, it was surprising that light cycle-restricted feeding caused a significant phase shift in the 24-hour rhythm of heart rate.(15) Light cycle-restricted feeding in mice also shifts the phase of circadian gene expression in peripheral tissues, including the heart, to align with the timing of feeding behavior.(17) Several studies implicate the cardiomyocyte circadian clock in regulating gene expression essential for cardiac excitability, cardiac electrophysiology, and heart rate.(8-12, 58, 60) One might predict that light cycle-restricted feeding shifts the phase of cardiomyocyte gene expression to facilitate changes in 24-hour rhythms of heart rate. However, genetically disrupting the cardiomyocyte-specific circadian clock (by inducing the deletion of *Bmal1*) did not abolish the light cycle-restricted feeding-mediated phase shift in the 24-hour rhythm in heart rate.(18) We conclude that autonomic receptor-mediated signaling largely shapes the 24-hour rhythm in heart rate during light cycle-restricted feeding.

### The 24-hour rhythm in the QT depends on core body temperature

We now report a robust linear QT-core body temperature relation in mice, and correcting the QT interval for core body temperature caused a loss in the 24-hour rhythm of the QT interval. These new data suggest the changes in the 24-hour rhythm of the QT interval in mice during light cycle-restricted feeding primarily reflect daily changes in core body temperature (not the RR interval). The idea that the 24-hour rhythm in the QT interval in mice primarily reflects core body temperature is also supported by a study showing that the 24-hour rhythm of the QT interval in mice is independent of the RR interval.(9) core body temperature was not reported in the study, but the observed change in the amplitude of the daily rhythm in the QT interval (4-5 ms) is consistent with a core body temperature amplitude typically seen in mice (1^°^C).(61)

The QT interval is used clinically in humans to assess the risk for certain deadly ventricular arrhythmias.(62) We extended our light cycle-restricted feeding studies to quantify changes in the QT interval in a transgenic knock-in mouse model that expresses a mutation linked to long QT syndrome in people.(33) Correcting the QT interval for core body temperature allowed direct comparisons of QT intervals at different core body temperatures across the 24-hour cycle, identified genotype-specific differences in the QT interval independent of core body temperature, and showed that light cycle-restricted feeding caused a prolongation of the QT interval in WT and *Scn5a*^+/ΔKPQ^ mice. These studies underscore the importance of monitoring and considering the influence of core body temperature when measuring QT intervals.

An important question is whether these data may be clinically relevant to humans. We propose that the daily rhythm in core body temperature also contributes to the 24-hour rhythm in the heart rate corrected QT (QTc) interval observed in people. Several studies report a temperature dependence of the QTc interval in people is 10-15 ms for every 1^°^C change in core body temperature.(24-29) The peak-to-trough amplitude in the daily core body temperature rhythm in people is about 1^°^C, and the peak-to-trough amplitude in the 24-hour rhythm in the QTc interval is 10-20 ms.(63, 64) Future studies that measure the QT interval and core body temperature across the 24-hour cycle in people will help delineate how much of the 24-hour rhythms in the QTc interval reflect temperature-dependent changes in the properties of ventricular repolarization.

Consideration of the influence of core body temperature on the QTc interval is warranted when the core body temperature might be different due to physiological, behavioral, environmental, or pharmacological changes (for example, patients with hypothermia or sepsis).

### Implications

Modern lifestyles disrupt evolutionarily conserved behavioral and environmental timekeeping cues critical for maintaining healthy circadian homeostatic mechanisms.(65) Our findings suggest that feeding behavior is a critical timekeeping cue for daily core body temperature and cardiac electrophysiology rhythms in mice. However, we did not see feeding as being a critical timekeeping cue for activity rhythms. This suggests that eating at an inverse time may cause misalignment between 24-hour rhythms in core body temperature, cardiac electrophysiology, and activity rhythms. If conserved, modern lifestyles that disrupt feeding rhythms in people may negatively impact the alignment between core body temperature, cardiac electrophysiological, and activity rhythms to impair cardiac performance and influence arrhythmogenic risk.

Another implication of this study is that it underscores the importance that daily rhythms in core body temperature have in shaping 24-hour rhythms in ventricular repolarization. Continuous core body temperature monitoring is not routinely performed in people undergoing electrophysiological testing despite its contribution to heart rate and QTc intervals. This is especially true in studies that monitor changes in heart rate and QTc intervals across the 24-hour cycle.

### Limitations

Observing a temperature dependence on the QT interval is not necessarily new and has been previously reported.(29) What is new is data suggesting the daily change in core body temperature is primarily responsible for generating a 24-hour rhythm in the QT interval. Ad libitum feeding may not be the best control for restricted light-cycle feeding. A dark cycle-restricted feeding control would rule out any potential effect of food consumption during the light phase seen in ad libitum-fed mice. Fasting mice makes them vulnerable to torpor (a transient hibernating-like state that causes a significant drop in heart rate and core body temperature below 32^°^C.(22) Our model of light cycle-restricted feeding did not calorically restrict the mice, and the mice continued to eat 3-4 grams of food per day during light cycle-restricted feeding (Figure 1S) and gained mass after one week (Figure 1S). Another limitation is that this study used only male mice. Results in female mice may differ. The mice used in the pharmacological study were older than those used in the initial light-cycle restricted feeding study. Whether a long QT_37°C_ interval is associated with an increased risk for spontaneous ventricular arrhythmias remains unclear. We found that, during light cycle-restricted feeding, the daily rhythm in the PR interval simultaneously shifted with the RR and QT intervals and core body temperature. However, due to the small absolute amplitude in the PR interval in response to light cycle-restricted feeding, it was difficult to determine if the PR interval reflected core body temperature, autonomic regulation of the heart, or both. The analyses in these studies focused on studying hourly mean values and 24-hour rhythms. We did not quantify the data at higher resolutions or study possible ultradian rhythms. Many cardiac electrophysiological channels that underlie the electrical rhythm of the heart, specifically ventricular repolarization, are not only temperature-sensitive but also time-of-day sensitive. Future studies examining the expression/activity of key components and circadian clock/autonomic nervous system markers in light cycle restricted feeding are warranted. Lastly, caution must be used when extrapolating data from mice to humans.(51, 66)

### Summary

Daily rhythms in cardiac electrophysiology align with feeding and activity cycles via signals originating from the SCN. Restricting food availability to the light cycle disrupts the association between environmental and behavioral timekeeping cues, resulting in the alignment of the daily rhythms of cardiac electrophysiology with the new feeding behavior. The alignment of the 24-hour rhythm in the RR interval primarily reflects changes in autonomic input to the heart. In contrast, the phase of the QT interval mirrors changes in core body temperature. Thermoneutrality studies suggest the distinct regulation of the 24-hour rhythms in RR and QT intervals also occurs in ad libitum-fed mice. We conclude that feeding-induced changes in daily autonomic signaling and core body temperature rhythms facilitate the alignment of 24-hour rhythms in heart rate and ventricular repolarization.

## Abbreviations

ECG: Electrocardiogram
pHF: high-frequency power
pLF: low-frequency power
SD: standard deviation
WT: Wild type
QT_37°C_: QT corrected to 37^°^C
ZT: Zeitgeber Time

## New and Noteworthy

We used time-restricted feeding and thermoneutrality to demonstrate that different mechanisms regulate the 24-hour rhythms in heart rate and ventricular repolarization. The daily rhythm in heart rate reflects changes in autonomic input, whereas daily rhythms in ventricular repolarization reflect changes in core body temperature. This novel finding has major implications for understanding 24-hour rhythms in mouse cardiac electrophysiology, arrhythmia susceptibility in transgenic mouse models, and interpretability of cardiac electrophysiological data acquired in thermoneutrality.

## Author contributions

All authors contributed to drafting the work and revising it critically for important intellectual content, approved the final version of the manuscript, and qualified for authorship.

## Data Availability

Data will be made available upon reasonable request.

## Acknowledgment

We want to thank Drs. Wayne Giles and Chao-Yin Chen (University of California, Davis) for providing feedback and discussion. We also acknowledge Isabel Stumpf (Centre College) and David Schneider (Centre College) for technical assistance.

