## Supplemental Figures for "Feeding Behavior Modifies the Circadian Variation in RR and QT intervals by Distinct Mechanisms in Mice"

### Slide 1
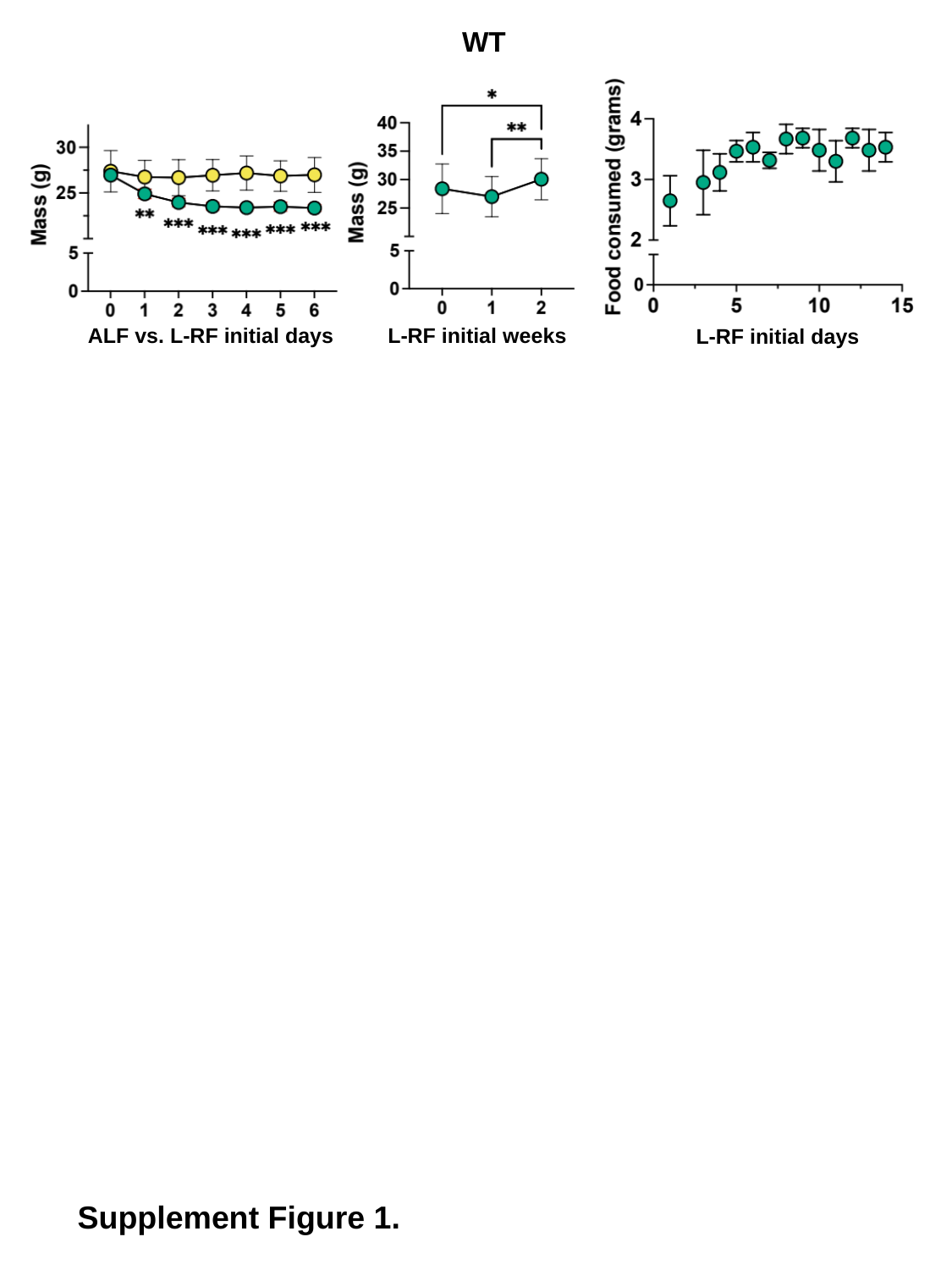

WT
ALF vs. L-RF initial days
L-RF initial weeks
L-RF initial days
Supplement Figure 1.

### Slide 2
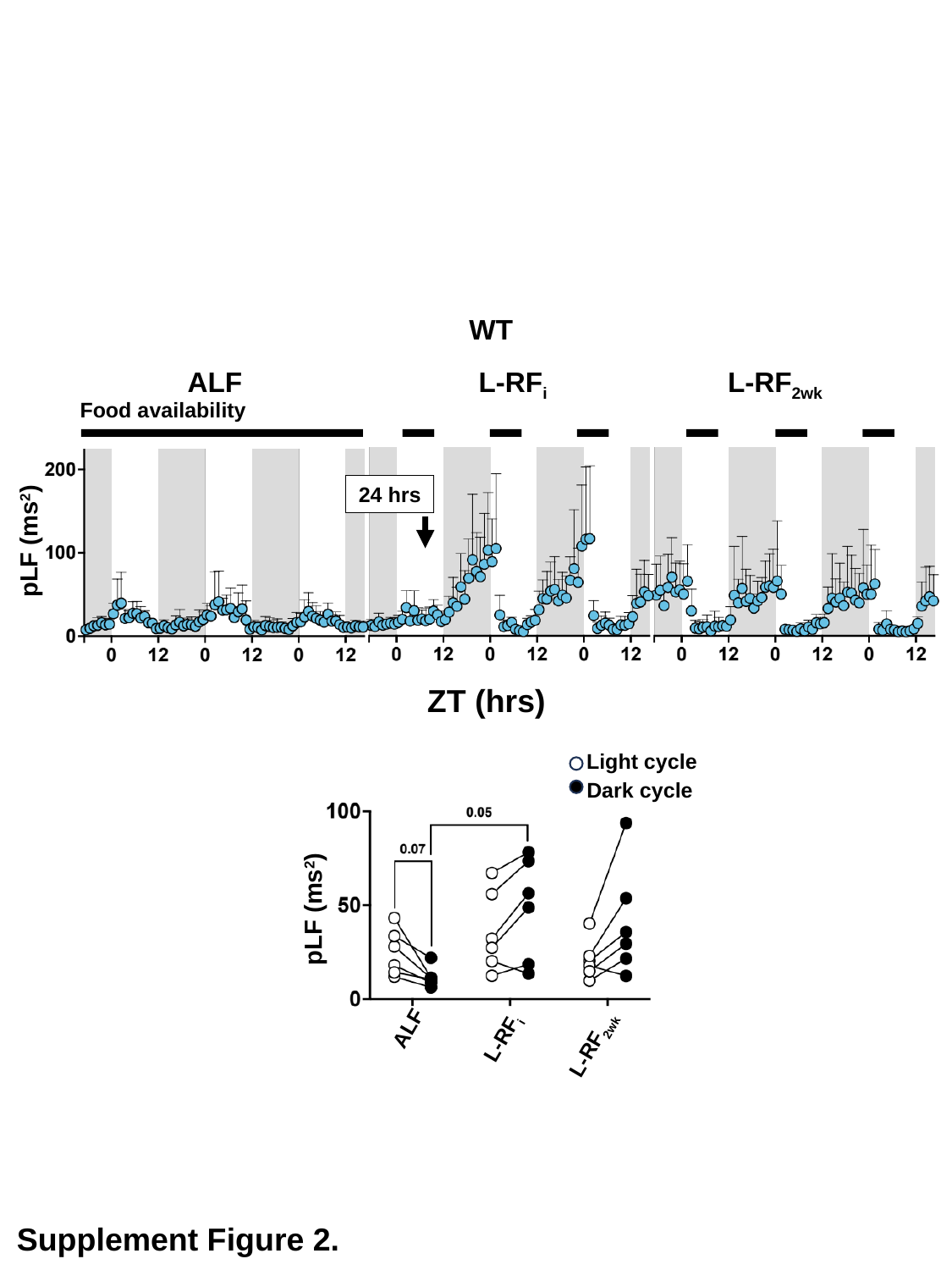

WT
ALF
L-RF_2wk_
L-RF_i_
Food availability
24 hrs
pLF (ms^2^)
ZT (hrs)
Light cycle
Dark cycle
pLF (ms^2^)
ALF
L-RF_i_
L-RF_2wk_
Supplement Figure 2.

### Slide 3
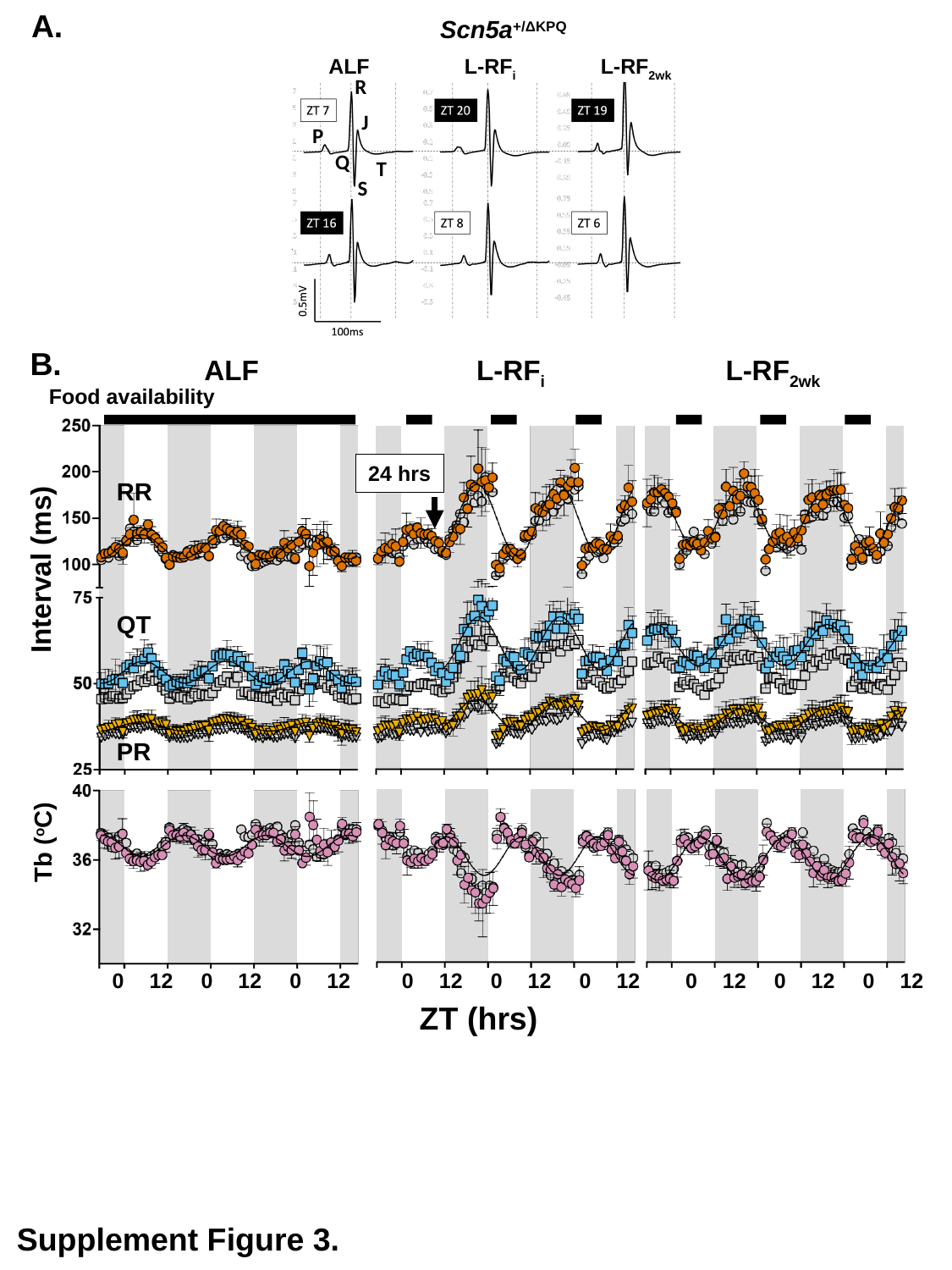

A.
Scn5a^+/ΔKPQ^
ALF
L-RF_i_
L-RF_2wk_
R
J
P
Q
T
S
B.
ALF
L-RF_2wk_
L-RF_i_
Food availability
24 hrs
RR
Interval (ms)
QT
PR
Tb (^o^C)
0
12
0
12
0
12
0
12
0
12
0
12
0
12
0
12
0
12
ZT (hrs)
Supplement Figure 3.

### Slide 4
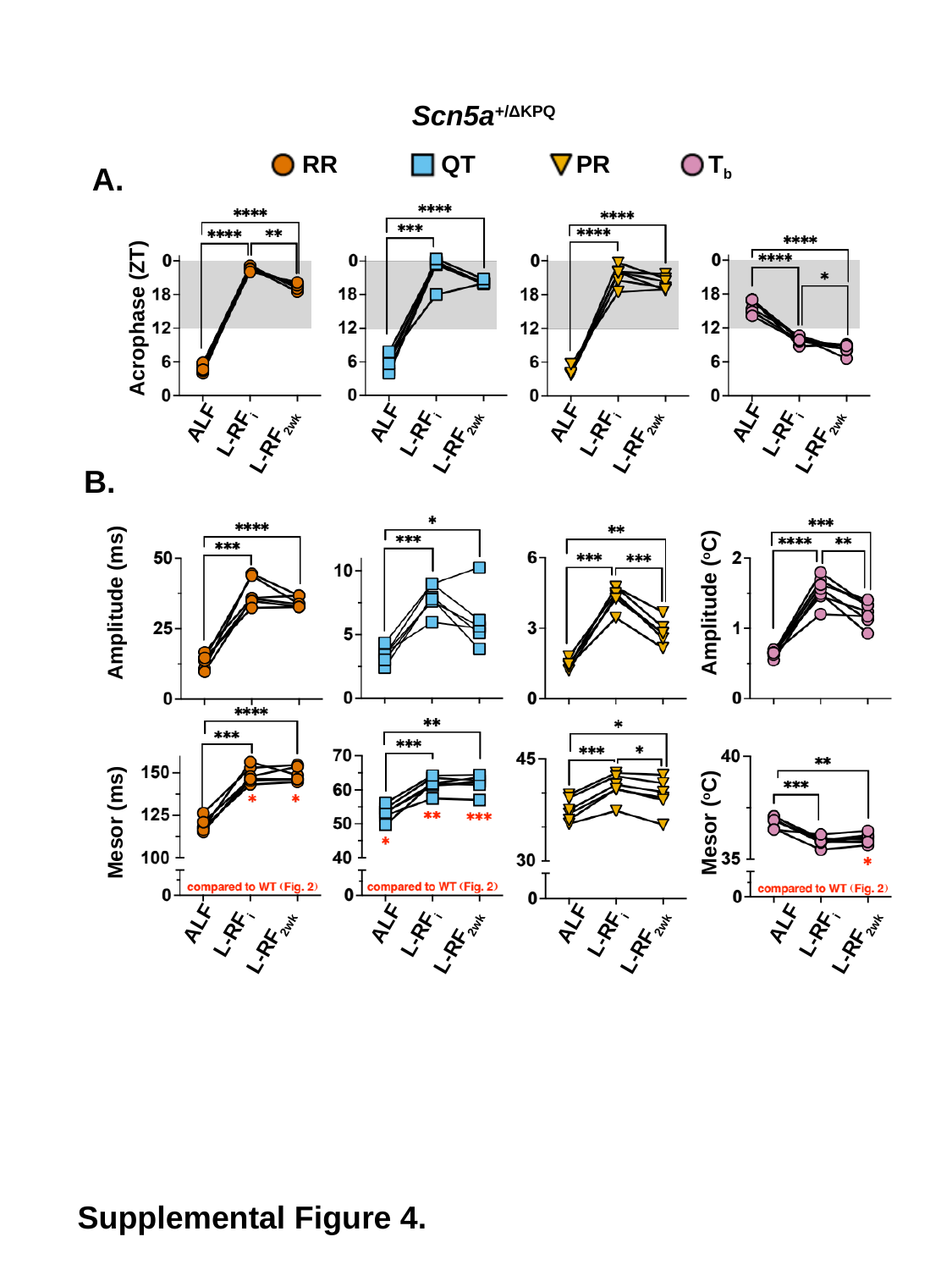

Scn5a^+/ΔKPQ^
RR
QT
PR
T_b_
A.
Acrophase (ZT)
ALF
ALF
ALF
ALF
L-RF_i_
L-RF_i_
L-RF_i_
L-RF_i_
L-RF_2wk_
L-RF_2wk_
L-RF_2wk_
L-RF_2wk_
B.
Amplitude (ms)
Amplitude (^o^C)
Mesor (ms)
Mesor (^o^C)
ALF
ALF
ALF
ALF
L-RF_i_
L-RF_i_
L-RF_i_
L-RF_i_
L-RF_2wk_
L-RF_2wk_
L-RF_2wk_
L-RF_2wk_
Supplemental Figure 4.

### Slide 5
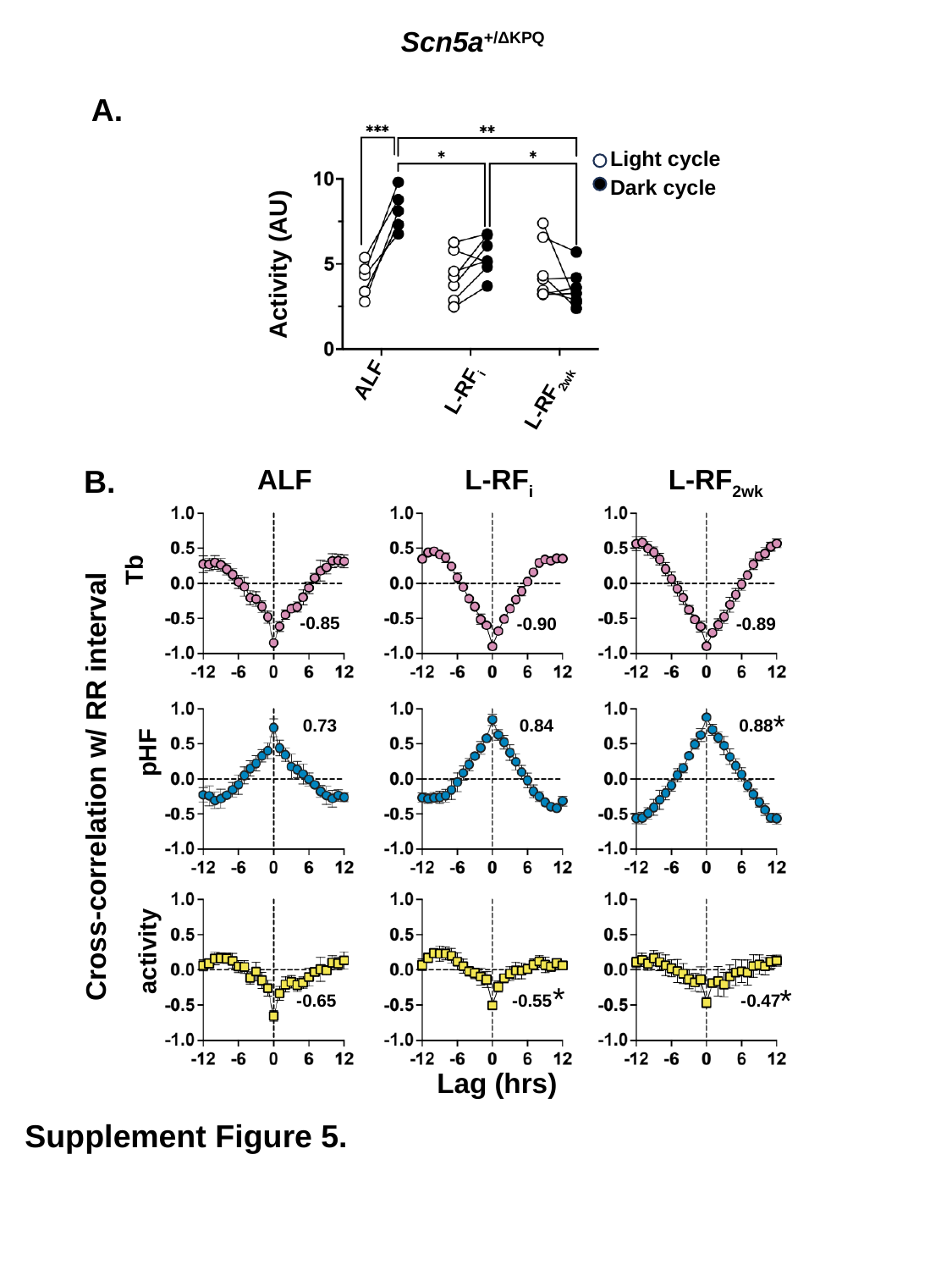

Scn5a^+/ΔKPQ^
A.
Light cycle
Dark cycle
Activity (AU)
ALF
L-RF_i_
L-RF_2wk_
B.
ALF
L-RF_i_
L-RF_2wk_
-0.85
-0.89
-0.90
*
0.73
0.84
0.88
pHF
Cross-correlation w/ RR interval
activity
*
*
-0.65
-0.55
-0.47
Lag (hrs)
Tb
Supplement Figure 5.
