## Supplementary material for "Feeding Behavior Modifies the Circadian Variation in RR and QT intervals by Distinct Mechanisms in Mice": legends

**Supplemental Figure Legends**

**Figure 1S. Light cycle-restricted feeding initially caused a decrease in mass, but the mice regained and increased mass after one week.** The left graph shows the daily body mass of mice in ALF (yellow) or after initially starting L-RF (blueish green, n = 5 mice per group, **p<0.01 ***p<0.001); the middle graph shows the average weekly mass in mice during the first 2 weeks of L-RF (n = 6, *p<0.05 and **p<0.01); and the right graph shows the average food consumption per mouse/day during each day after the start of L-RF for two weeks (n=6).

**Figure 2S. Light cycle-restricted feeding and its impact on heart rate variability measured in the low-frequency domain is shown. A.** The graphs show the corresponding data for the power of the low-frequency component in heart rate variability (pLF, sky blue). The left graph shows the data from mice in ad libitum-conditions (ALF), the middle graph shows the data initially starting light cycle-restricted feeding (L-RF_i_), and the right graph shows the data two weeks after starting light cycle-restricted feeding (L-RF_2wk_). Data were plotted as a function of ZT and the shaded regions correspond to the dark cycle (n = 6 mice, data points are averaged data and SD). **B.** The graph shows the average pLF value for each mouse during the light cycle (open circle) and dark cycle (solid circle) measured during ALF, L-RF_i_, and L-RF_2wk_. Data were analyzed using a one-way ANOVA with Geisser and Greenhouse correction and Sidǎk correction, n = 6 mice per group.

**Supplemental Table 1.** Results from the cosine waveform fit to the data recorded from individual mice. Included are the JTK p-values. RR interval data measured from mouse WT 17 two weeks after the start of light cycle-restricted feeding had a JTK p value > 0.05 and were included in cosine analyses.
